# Investigating the eukaryotic host-like SLiMs in microbial mimitopes and their potential as novel drug targets for treating autoimmune diseases

**DOI:** 10.1101/2022.06.20.496681

**Authors:** Anjali Garg, Neelja Singhal, Manish Kumar

## Abstract

Several pathogens sustain themselves inside the host by mimicking short linear motifs (SLiMs) of the host proteins. SLiMs are short stretches of 3-10 amino acids which are functionally diverse and mediate various signaling and protein-protein interactions (PPIs). Hence, mimicry of the host- SLiMs helps the microbes in modulating/disrupting the host defense mechanisms. This is the first report investigating the evolutionary characteristics and presence of eukaryotic host-like SLiMs in microbial peptides (mimitopes). Evaluation of the selection pressure revealed that 60% of the bacterial and 25% of the viral mimitopes which overlapped with the host-like SLiMs were evolutionarily conserved (*ω* < 1). Interestingly, host-like SLiMs were abundant in mimicry proteins but were less frequent in microbial mimitopes. This reflects that the majority of the pathogens cannot potentially rewire the host PPI networks for their advantage, but some can. Of the 152 bacterial and 43 viral mimitopes investigated only 10 bacterial and 4 viral mimitopes showed SLiMs. This indicates that mimitopes of some pathogens can be explored as novel drug targets for eliminating the etiopathological agent and treating the autoimmune disease, thereof. The repertoire of mimitopes identified here might provide important clues for the discovery of new drugs/protein-based immune-modulatory molecules against the pathogens.

**Key points:**

1. Mimicry of the host- SLiMs helps the microbes in modulating/disrupting the host defense mechanisms.
2. Host-like SLiMs were abundant in mimicry proteins but were less frequent in microbial mimitopes.
3. Evaluation of the selection pressure revealed that 60% of the bacterial and 25% of the viral mimitopes which overlapped with the host-like SLiMs were evolutionarily conserved.

## Introduction

The host genetics play an important role in the induction of autoimmune response against self-antigens however; several epidemiological and molecular evidences suggest microbial pathogens (viruses and bacteria) are the principal environmental triggers of autoimmunity [1–5]. Reportedly, four main criteria have been associated with molecular mimicry and the underlying human disease, (i) epidemiological evidence associating a pathogen with a disease, (ii) presence of antibodies and/or immune cells against the specific antigens related with the disease, (iii) cross-reactivity of antibodies or immune cells of the host with microbial antigens and, (iv) reproduction by the antigen of the disease process *in vivo* or *in vitro* [6,7].

Microbes can exhibit four types of molecular mimicry with the host proteins, (i) sequential or structural similarities with the proteins or protein-domains, (ii) similarities with the protein-structures without sequence homology, (iii) architectural similarities with the binding surfaces without sequence homology (interface mimicry) and, (iv) similarities with the short linear motifs (SLiMs) of the proteins (motif mimicry) [8,9]. SLiMs are short stretches of amino acids (3-10 amino acids) which are functionally diverse and mediate various signaling and protein-protein interactions (PPIs) [10,11]. Reportedly, several viruses and bacteria propagate and sustain themselves inside the host by mimicking SLiMs of the host proteins [12].

PPIs and cell signaling play a pivotal role in the functioning of cellular life. Hence, mimicry of the host proteins involved in PPIs and cell signaling greatly helps microbes in modulating/disrupting the host defense mechanisms. Moreover, substitution of microbial proteins in the host PPI networks eventually shifts the paradigm of host-pathogen interactions in the favor of the pathogen. Several eukaryotic host-like SLiMs have been reported in viral proteins which helps in their entry inside the host cell and modulation of host cellular pathways related to transcription regulation, cell cycle, immune response etc [13–15]. Similarly, several eukaryotic host-like SLiMs have also been reported in bacterial proteins which helps them to interfere with signaling pathways involving protein tyrosine kinases (TKs) and mitogen-activated protein kinases (MAPKs), tyrosine phosphorylation etc [12,16–19]. The emergence of antibiotic resistance in pathogens necessitates discovery of new anti-infective therapies. Recently, protein-based immune-modulatory molecules or drugs which can disrupt the host-pathogen PPIs have been proposed as a novel anti-infective therapy [12]. Such therapeutics can interfere with the host-pathogen SLiM-mediated interactions and create a hostile and non-conducive host environment for the survival of the pathogen. Though, mimicry of the eukaryotic host-like SLiMs has been recognized as an important mechanism underlying microbial pathogenicity, presence and characteristics of host-like SLiMs have not been studied in the mimicry peptides (mimitopes) of bacteria and viruses experimentally associated with autoimmune diseases. Thus, the present study was conducted to discern if the experimentally verified microbial mimitopes underlying various autoimmune diseases exhibit motif mimicry with the host and potentially modulate the host PPIs. Additionally, the evolutionary pressure on microbial mimitopes was also determined. This is the first report on potential of autoimmunity-related microbial mimitopes in modulating host protein-protein interactions and their evolutionary characteristics.

## Material and Methods

### Retrieval of experimentally validated mimicry proteins from miPepBase

In the present study, the information on bacterial, viral and the host mimicry proteins was retrieved from a database of experimentally verified mimicry proteins, miPepBase (updated version assessed on January 2021 [26]. The database collates information about only the experimentally verified mimicry proteins/peptides and autoimmune diseases, thereof [20].

### Investigating the presence of eukaryotic host-like SLiMs in microbial mimicry proteins and mimitopes

The presence of eukaryotic host-like SLiMs in the microbial mimicry proteins and mimitopes was predicted using ANCHOR [21]. A eukaryotic host-like SLiM was considered to be present in the microbial mimitope if at least half of the amino acid of mimitopes overlapped with the SLiM region.

### Molecular evolutionary analyses of microbial mimitopes

The putative homologous/orthologous sequence of microbial mimicry proteins were identified using BLAST search against NCBI non-redundant (NR) protein database. To obtain a sequence homologous to each microbial protein, the NR database was searched using the microbial protein as query. Search results with query coverage of ≥80 and atleast 80% sequence identity were selected as the criteria for homology and used for further evolutionary analyses. The Orthologous protein clusters of each mimicry protein were aligned using ClustalW version 2 [22]. The corresponding gene sequences of these proteins were also aligned on the basis of their codons using pal2nal [23]. The ratio between the rate of non-synonymous substitution to the rate of synonymous substitution (*ω* = *d*N/*d*S) was used as a measure of the strength of selection pressure acting on a protein-coding gene. Assuming synonymous mutations are subjected to almost strictly neutral selection, the *ω* < 1, *ω* = 1, and *ω* > 1 represent negative selection, neutral evolution, and positive Darwinian selection, respectively [24]. To compute the dN/dS ratio for each amino acid of the microbial mimitope, a site-specific model of the likelihood method was used using the codeml module of the PAML package [25].

## Results and Discussion

Microbial mimicry of the host peptides has been implicated in several autoimmune diseases like multiple sclerosis [26], type 1 diabetes mellitus [27], autoimmune uveitis [28], encephalomyelitis [29], inflammatory bowel disease [30,31], Crohn’s disease [32,33], sarcoidosis, etc. Motif mimicry of the eukaryotic host-like SLiMs has been associated with the pathogenicity of several bacteria and viruses because it helps them to hijack the host processes by rewiring the host PPI networks, signaling pathways, post-translational modifications etc [10,11]. Since, SLiMs are abundantly present in cell signaling and PPI network proteins, drugs targeting SLiM-regulated cellular processes have been proposed as novel therapeutics against bacterial and viral infections [12,13,34]. Thus, the present study was conducted to discern if the mimicry peptides (experimentally verified as responsible for autoimmune diseases; collated in miPepBase) contain the functional modules of PPIs, SLiMs and can be explored as novel drug targets.

A total of 147 bacterial mimicry proteins with their corresponding 27 host proteins and, 34 viral mimicry proteins with their corresponding 22 host mimicry proteins were retrieved from the miPepBase. The bacterial and their corresponding host mimicry proteins were named as Bacterial-set proteins. The viral and their corresponding host mimicry proteins were named as Viral-set proteins. The Bacterial-set proteins were involved in 16, while the Viral-set proteins were involved in 12 different types of autoimmune diseases. The detailed information of the host and pathogen, UniProtKB ID and name of the mimicry-protein(s), mimitope sequences, associated autoimmune diseases, and source of information (Pubmed ID) is provided in Table S1.

During infection, the pathogens repurpose the host cells in a manner which is conducive to their own survival, proliferation and helpful for evading the host immunity. Thus, if the host cells can be rewired to the uninfected state or the one that is close to it, the disease progression can be halted or slowed down. Thus, drugs/molecules which can inhibit either (i) interactions between microbial host-like SLiMs and host PPI networks or, (ii) the metabolic chokepoints of these PPI networks or, (iii) nodes in the regulatory networks upstream or downstream of the hijacked host-SLiMs can be explored as novel therapeutics. Also termed as host-directed therapy, this approach has proved useful in combating many viruses and bacteria [12]. The classical examples of host-directed therapy are the chemical inhibition of the proteasomal pathway which resulted in reduced viral loads in dengue infection and, RNA interference (RNAi) against some human proteins which inhibited replication of HIV and hepatitis C virus [35]. Our results revealed that the eukaryotic host-like SLiMs were abundant in the microbial mimicry proteins because 41 bacterial and 20 viral mimicry proteins showed the presence of SLiMs (Table S2). On the contrary, only a few microbial mimitopes overlapped with the eukaryotic host-like SLiMs. Of the total 152 bacterial mimitopes, 10 bacterial mimitopes showed SLiMs and of the 43 viral mimitopes only 4 viral mimitopes had SLiMs (Table 1). This suggests that the most of the microbial mimitopes implicated in autoimmune diseases could not potentially rewire the host PPI networks for their own benefit. And, their pathology might have only the autoimmune component due to cross reactivity between the microbial epitopes and host peptides. However, the microbial mimitopes of ten bacteria and three viruses overlapped with the eukaryotic host-like SliMs (Table 1). Seven mimitopes, one each of *Agrobacterium tumefaciens, Yersinia enterocolitica, Escherichia coli, Propionibacterium freudenreichii, Streptococcus mutans, Lactobacillus johnsonii, Bifidobacterium longum* and two each of *Mycobacterium leprae* and Group A streptococcus showed the presence of eukaryotic host-like SLiMs in their mimitopes. Among the viruses, one mimitope of Herpes simplex virus and Herpes virus saimiri and two mimitopes of Epstein Barr virus had host-like SLiMs (Table 1). This suggests that these peptide regions might not only be responsible for inducing autoimmune diseases in the host by exhibiting epitope mimicry with the host, motif mimicry of the host-like SLiMs might also aid these bacteria and viruses in interrupting the host cell signaling and PPI networks.

**Table 1:**
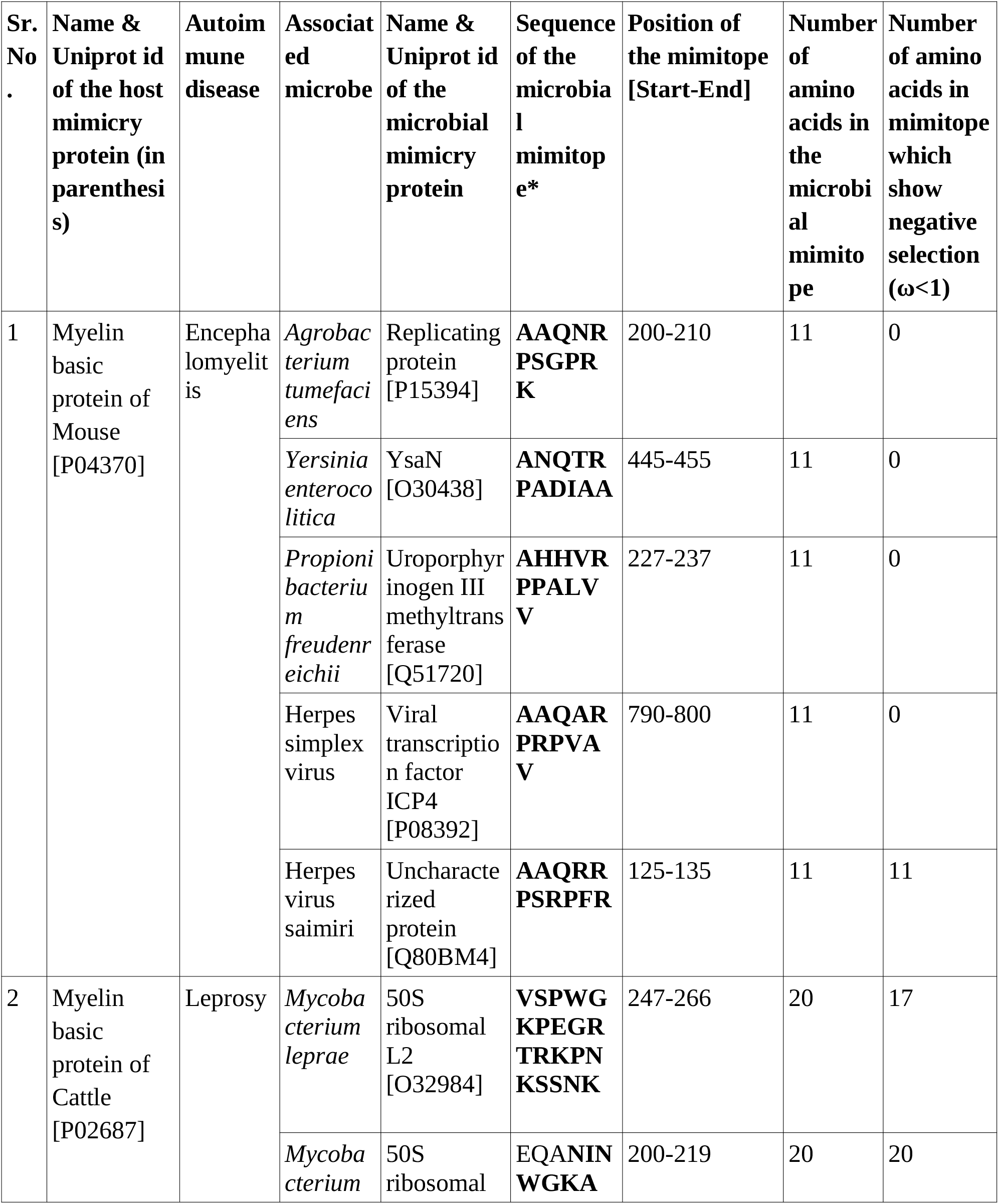
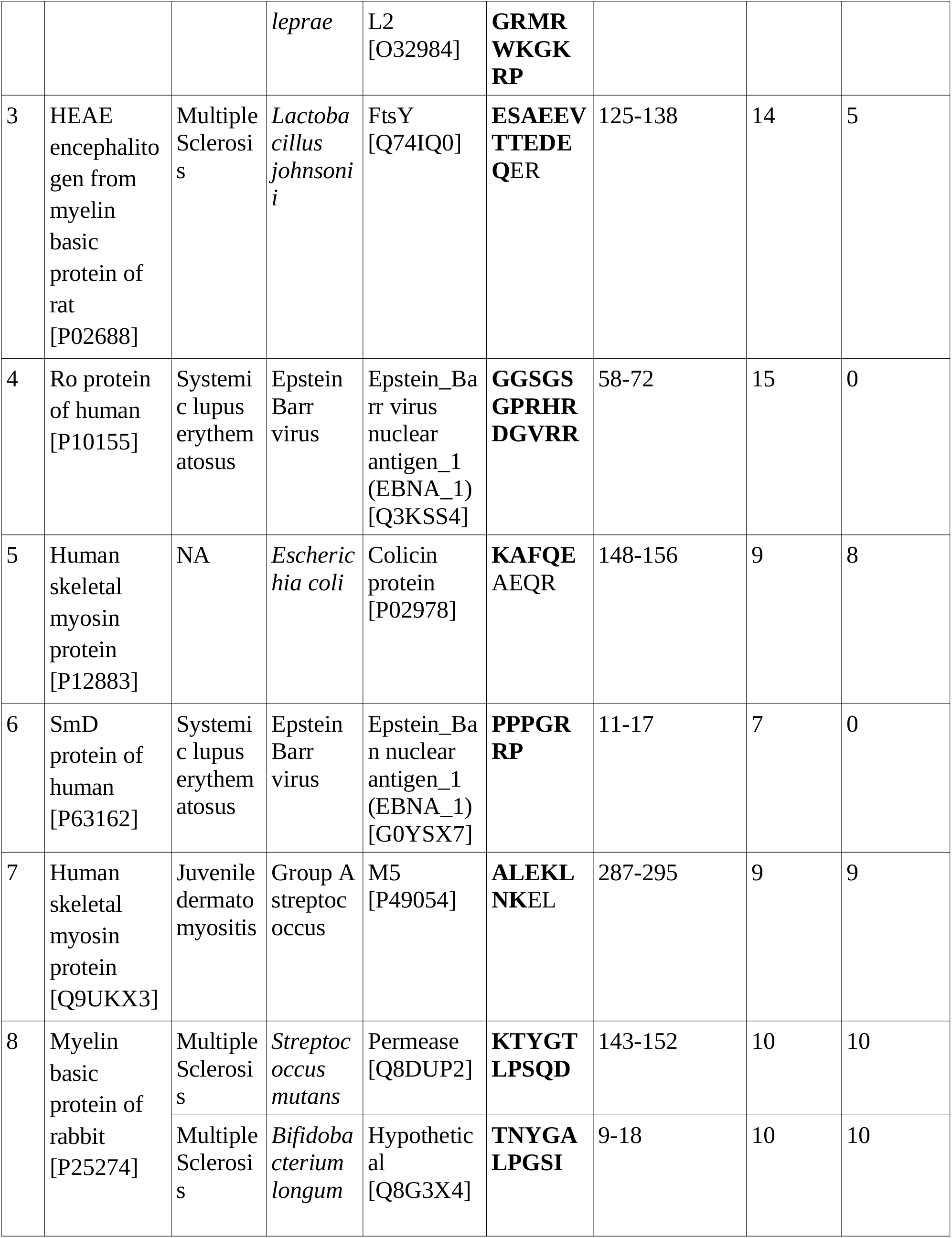
Detailed information about the microbial mimitopes which overlapped with the eukaryotic host-like SliMs. SLiM regions in the microbial mimitopes are shown in bold face.

Evaluation of the selection pressure on the microbial mimitopes revealed that 77.85% of the bacterial and 83.54% of the viral amino acids were under negative selection pressure (*ω* < 1), implying that microbial mimtopes were generally conserved or had a low mutation rate. Comas et al., reported that the T cell epitopes of *M. tuberculosis* were highly conserved which might be a distinct evolutionary strategy of a highly successful pathogen as *M. tuberculosis* for immune subversion [36]. This suggests that other pathogens might also employ a similar strategy for a sustained survival inside the host. However, analysis of the microbial mimitopes which overlapped with the eukaryotic host-like SLiMs revealed that only 40% of the bacterial mimitopes and 25% of the viral mimitopes were under negative selection pressure (*ω* < 1) (Table 1). This observation can be explained by the fact that SLiMs are disordered regions of the proteins hence, the rate of mutation is high and these regions are usually under a positive selection pressure [37]

Conventionally, autoimmune diseases like systemic lupus erythematosus, encephalomyelitis, multiple sclerosis etc. are treated using immunomodulators or immunosuppressants which only try to cure the symptoms but do not eliminate the etiological agent which continues to proliferate inside the host and interfering with vital cellular process(es). On the contrary, protein-based immune-modulatory molecules/drug molecules designed to inhibit the mimitopes (or eukaryotic host-like SLiMs) might also disrupt the host-pathogen PPIs. This implies that inhibitors of microbial mimitopes (or eukaryotic host-like SLiMs) identified in this study would not only help in treating the pathogen-associated autoimmune disease but also eliminate the etiopathological agent associated with the disease.

Finally, the repertoire of mimitopes discovered in this study suggests new paradigms underlying autoimmune diseases and the etiopathology associated with the infection. Such knowledge can provide important clues for the discovery of new drugs/ protein-based immune-modulatory molecules against the pathogens.

## Acknowledgments

The work was carried out using the resources funded by the Science and Engineering Research Board under Fast Track Proposals for Young Scientists Scheme [Grant No: SR/FT/LS-84/2010] and UGC Major Research Project [Grant No: 41-38/2012(SR)]. AG is supported by ICMR-JRF scheme [Grant Number: 3/1/3 J.R.F.-2016/LS/HRD-(32262)]. NS is supported by CSIR Senior Research Associateship (Scientists’ Pool Scheme) [Grant Number: 13(9089-A)/2019-Pool].

## Author contributions

NS and MK conceived and designed the study. AG performed the work, prepared the illustrations and wrote initial draft manuscript. All authors contributed to the critical analysis and revision. All authors analyzed the results and wrote the final versions of manuscript.

## Figure legends

**Figure 1:**
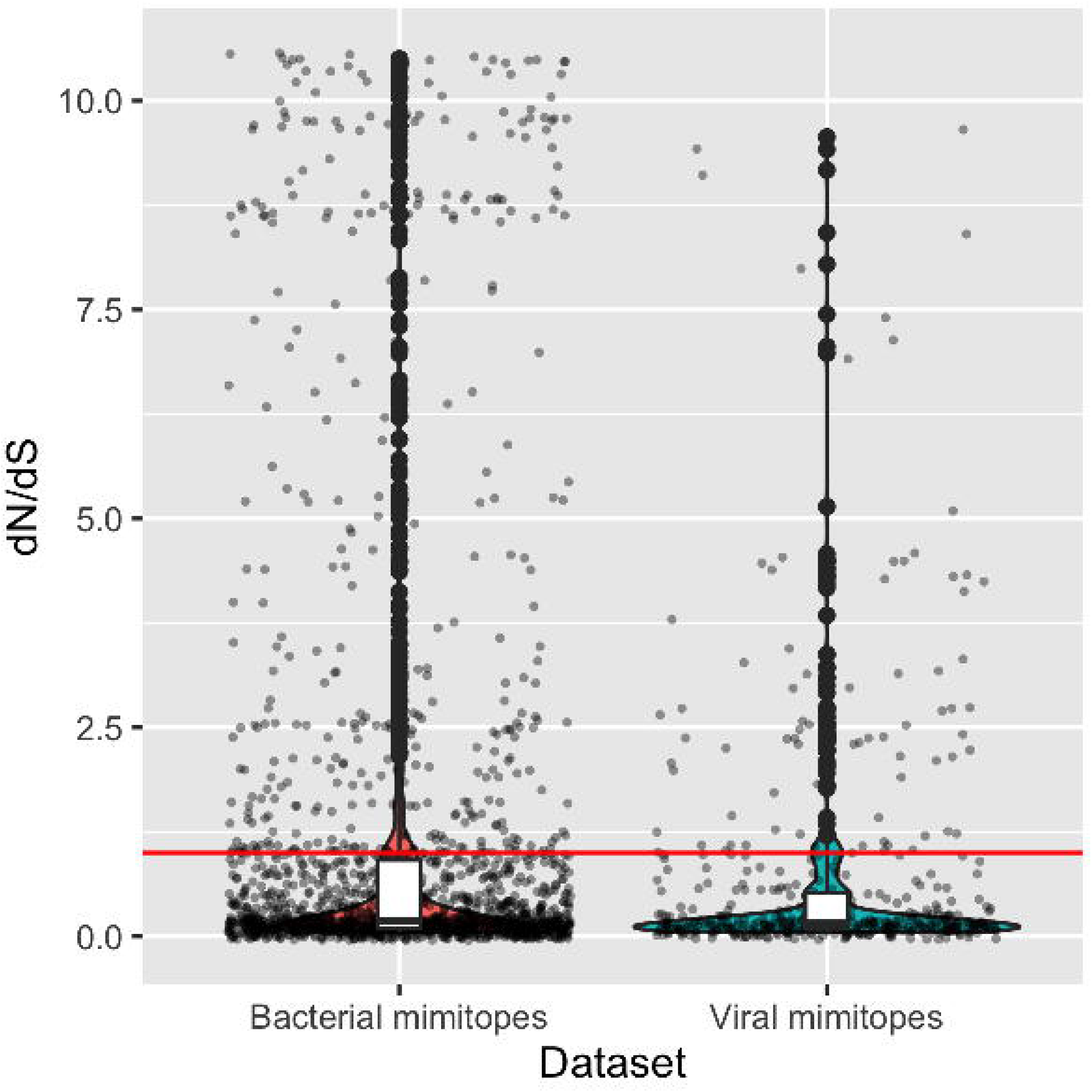
Distribution of site-specific omega values in bacterial and viral mimitopes. The horizontal red line represents dN/dS =1. Amino acids with omega value below 1 are under purifying/negative selection and residues above the line are under positive selection.

## Supplementary tables

**Table S1:** Detailed information about mimicry proteins, peptides and the associated autoimmune diseases of bacteria, viruses and hosts. The data was retrieved from miPepBase database (accessed on January 2021).

**Table S2:** Detailed information about the SLiMs in the bacterial and viral mimicry proteins.

## Reference

1. Fujinami RS. Viruses and autoimmune disease – two sides of the same coin? Trends Microbiol. 2001; 9:377–381

2. Libbey JE, Fujinami RS. Potential triggers of MS. Results Probl. Cell Differ. 2010; 51:21–42

3. Ascherio A, Munger KL. Environmental risk factors for multiple sclerosis. Part II: Noninfectious factors. Ann. Neurol. 2007; 61:504–513

4. Tarakhovsky A, Prinjha RK. Drawing on disorder: How viruses use histone mimicry to their advantage. J. Exp. Med. 2018; 215:1777–1787

5. Lester PJ, Buick KH, Baty JW, et al. Different bacterial and viral pathogens trigger distinct immune responses in a globally invasive ant. Sci. Rep. 2019; 9:5780

6. Ang CW, Jacobs BC, Laman JD. The Guillain-Barré syndrome: a true case of molecular mimicry. Trends Immunol. 2004; 25:61–66

7. Johnson TP, Tyagi R, Lee PR, et al. Nodding syndrome may be an autoimmune reaction to the parasitic worm Onchocerca volvulus. Science Translational Medicine 2017; 9:eaaf6953

8. Xue B, Blocquel D, Habchi J, et al. Structural disorder in viral proteins. Chem. Rev. 2014; 114:6880–6911

9. Dolan PT, Roth AP, Xue B, et al. Intrinsic disorder mediates hepatitis C virus core-host cell protein interactions: HCV Core MoRFs Target Cellular Proteins. Protein Sci. 2015; 24:221–235

10. Davey NE, Van Roey K, Weatheritt RJ, et al. Attributes of short linear motifs. Mol. Biosyst. 2012; 8:268–281

11. Dinkel H, Van Roey K, Michael S, et al. The eukaryotic linear motif resource ELM: 10 years and counting. Nucleic Acids Res. 2014; 42:D259–66

12. Sámano-Sánchez H, Gibson TJ. Mimicry of Short Linear Motifs by Bacterial Pathogens: A Drugging Opportunity. Trends Biochem. Sci. 2020; 45:526–544

13. Davey NE, Travé G, Gibson TJ. How viruses hijack cell regulation. Trends in Biochemical Sciences 2011; 36:159–169

14. Garamszegi S, Franzosa EA, Xia Y. Signatures of Pleiotropy, Economy and Convergent Evolution in a Domain-Resolved Map of Human–Virus Protein–Protein Interaction Networks. PLoS Pathogens 2013; 9:e1003778

15. Hagai T, Azia A, Babu MM, et al. Use of host-like peptide motifs in viral proteins is a prevalent strategy in host-virus interactions. Cell Rep. 2014; 7:1729–1739

16. Zhu Y, Li H, Long C, et al. Structural insights into the enzymatic mechanism of the pathogenic MAPK phosphothreonine lyase. Mol. Cell 2007; 28:899–913

17. Li H, Xu H, Zhou Y, et al. The Phosphothreonine Lyase Activity of a Bacterial Type III Effector Family. Science 2007; 315:1000–1003

18. Higashi H, Tsutsumi R, Fujita A, et al. Biological activity of the Helicobacter pylori virulence factor CagA is determined by variation in the tyrosine phosphorylation sites. Proc. Natl. Acad. Sci. U. S. A. 2002; 99:14428–14433

19. Frese S, Schubert W-D, Findeis AC, et al. The phosphotyrosine peptide binding specificity of Nck1 and Nck2 Src homology 2 domains. J. Biol. Chem. 2006; 281:18236–18245

20. Garg A, Kumari B, Kumar R, et al. miPepBase: A Database of Experimentally Verified Peptides Involved in Molecular Mimicry. Front. Microbiol. 2017; 8:2053

21. Dosztányi Z, Mészáros B, Simon I. ANCHOR: web server for predicting protein binding regions in disordered proteins. Bioinformatics 2009; 25:2745–2746

22. Larkin MA, Blackshields G, Brown NP, et al. Clustal W and Clustal X version 2.0. Bioinformatics 2007; 23:|p|

23. Suyama M, Torrents D, Bork P. PAL2NAL: robust conversion of protein sequence alignments into the corresponding codon alignments. Nucleic Acids Res. 2006; 34:W609–12

24. Yang Z. Inference of selection from multiple species alignments. Curr. Opin. Genet. Dev. 2002; 12:688–694

25. Yang Z. PAML 4: phylogenetic analysis by maximum likelihood. Mol. Biol. Evol. 2007; 24:1586–1591

26. Wekerle H, Hohlfeld R. Molecular Mimicry in Multiple Sclerosis. New England Journal of Medicine 2003; 349:185–186

27. Coppieters KT, Wiberg A, von Herrath MG. Viral infections and molecular mimicry in type 1 diabetes. APMIS 2012; 120:941–949

28. Wildner G, Diedrichs-Möhring M. Autoimmune uveitis induced by molecular mimicry of peptides from rotavirus, bovine casein and retinal S-antigen. Eur. J. Immunol. 2003; 33:2577–2587

29. Phelan J, Grabowska AD, Sepúlveda N. A potential antigenic mimicry between viral and human proteins linking Myalgic Encephalomyelitis/Chronic Fatigue Syndrome (ME/CFS) with autoimmunity: The case of HPV immunization. Autoimmun. Rev. 2020; 19:102487

30. Yusung S, Braun J. Molecular mimicry, inflammatory bowel disease, and the vaccine safety debate. BMC Med. 2014; 12:166

31. Biank V, Broeckel U, Kugathasan S. Pediatric inflammatory bowel disease: Clinical and molecular genetics. Inflammatory Bowel Diseases 2007; 13:1430–1438

32. Polymeros D, Tsiamoulos ZP, Koutsoumpas AL, et al. Bioinformatic and immunological analysis reveals lack of support for measles virus related mimicry in Crohn’s disease. BMC Med. 2014; 12:139

33. Cunha-Neto E, Kalil J. Molecular Mimicry and Chagas’ Disease. Molecular Mimicry, Microbes, and Autoimmunity 2014; 257–274

34. Via A, Uyar B, Brun C, et al. How pathogens use linear motifs to perturb host cell networks. Trends Biochem. Sci. 2015; 40:36–48

35. Schwegmann A, Brombacher F. Host-Directed Drug Targeting of Factors Hijacked by Pathogens. Sci. Signal. 2008; 1:re8–re8

36. Comas I, Chakravartti J, Small PM, et al. Human T cell epitopes of Mycobacterium tuberculosis are evolutionarily hyperconserved. Nature Genetics 2010; 42:498–503

37. Uyar B, Weatheritt RJ, Dinkel H, et al. Proteome-wide analysis of human disease mutations in short linear motifs: neglected players in cancer? Mol. Biosyst. 2014; 10:2626–2642

